# Deficiency of macrophage-derived Dnase1L3 causes lupus-like phenotypes in mice

**DOI:** 10.1101/2023.04.17.537232

**Authors:** Minal Engavale, Colton J. Hernandez, Angelica Infante, Tanya LeRoith, Elliott Radovan, Lauryn Evans, Johanna Villarreal, Christopher M. Reilly, R. Bryan Sutton, Peter A. Keyel

## Abstract

Systemic Lupus Erythematosus (SLE) is a chronic autoimmune disease caused by environmental factors and loss of key proteins. One such protein is a serum endonuclease secreted by macrophages and dendritic cells, Dnase1L3. Loss of Dnase1L3 causes pediatric-onset lupus in humans is Dnase1L3. Reduction in Dnase1L3 activity occurs in adult-onset human SLE. However, the amount of Dnase1L3 necessary to prevent lupus onset, if the impact is continuous or requires a threshold, and which phenotypes are most impacted by Dnase1L3 remain unknown. To reduce Dnase1L3 protein levels, we developed a genetic mouse model with reduced Dnase1L3 activity by deleting *Dnase1L3* from macrophages (cKO). Serum Dnase1L3 levels were reduced 67%, though Dnase1 activity remained constant. Sera were collected weekly from cKO and littermate controls until 50 weeks of age. Homogeneous and peripheral anti-nuclear antibodies were detected by immunofluorescence, consistent with anti-dsDNA antibodies. Total IgM, total IgG, and anti-dsDNA antibody levels increased in cKO mice with increasing age. In contrast to global Dnase1L3^−/−^ mice, anti-dsDNA antibodies were not elevated until 30 weeks of age. The cKO mice had minimal kidney pathology, except for deposition of immune complexes and C3. Based on these findings, we conclude that an intermediate reduction in serum Dnase1L3 causes mild lupus phenotypes. This suggest that macrophage-derived DnaselL3 is critical to limiting lupus.

## Introduction

Systemic Lupus Erythematosus (SLE) is a challenging autoimmune disease to treat due to the environmental and genetic heterogeneity of the disease. In order to understand polygenic SLE disease pathogenesis, we first need to understand how individual genes contribute to disease phenotypes in SLE. The key to unraveling the genetic heterogeneity of SLE are mutations in single genes that cause monogenic SLE. One gene associated with monogenic SLE in both humans and in mice is the serum endonuclease *Dnase1L3* [1, 2]. *Dnase1L3* loss in humans causes familial, pediatric-onset lupus with 100% penetrance, average onset by age 6, no gender bias, high (∼64%) incidence of lupus nephritis, and high anti-nuclear and anti-dsDNA antibodies [1]. Partial loss of Dnase1L3 via activity-reducing mutations causes the related disease hypocomplementemic urticarial vasculitis syndrome (HUVS) [3, 4]. HUVS is characterized by decreased C1q, serum complement activation, arthritis, urticarial vasculitis, no anti-dsDNA antibodies, and patients may die from pulmonary obstruction [5]. Reduced Dnase1L3 serum activity occurs in human adult-onset SLE [2, 6, 7], in part due to neutralizing autoantibodies targeting Dnase1L3 [8, 9]. Dnase1L3 is secreted primarily by macrophages and dendritic cells (DCs) [2, 10], with IL-4 polarized macrophages secreting more Dnase1L3 [11]. Dnase1L3 is upstream of T/B cell responses, inflammation, and complement. Thus, reduced Dnase1L3 activity contributes to SLE disease heterogeneity, and pathogenesis.

Understanding the pathogenesis caused by loss of Dnase1L3 activity is complicated by the presence of a second serum endonuclease, Dnase1. Both endonucleases are part of the Dnase1 family, and degrade cell-free DNA in the serum [12]. While Dnase1 has been implicated in lupus-like phenotypes in mice [13], recent work showed that B6 mice lacking Dnase1 do not develop lupus-like phenotypes, nor does loss of Dnase1 exacerbate lupus-like phenotypes in Dnase1L3^−/−^ mice [14]. One key difference between Dnase1 and Dnase1L3 is a disordered C-terminus that enables digestion of DNA complexed with proteins and lipids [2, 15, 16]. While both nucleases can degrade plasmid DNA, Dnase1L3 has specific activity against complexed DNA that is essential for preventing SLE.

Mice are a powerful model system to better understand disease heterogeneity in autoimmunity. The polygenic lupus mouse models NZB/W F1 and MRL/*lpr* express a mutant Dnase1L3 with reduced activity [17]. Lupus-like phenotypes arise in strains not prone to autoimmunity (i.e. pure C57Bl/6 background) when Dnase1L3^−/−^ is globally eliminated [2, 10]. Global Dnase1L3^−/−^ mice produce anti-dsDNA antibodies in the first two months of life, and expand B cells and neutrophils expand in the first 8-14 weeks [2, 10, 14]. However, in global Dnase1L3^−/−^ mice, the full repertoire of autoantibodies does not develop until ∼50 weeks [2]. By 50 weeks of age, Dnase1L3^−/−^ mice develop splenomegaly with spontaneous germinal center formation, glomerulonephritis, and kidney IgG deposition [2, 18]. Pathological changes in the kidney were only mildly evident [2]. Thus, mice can be used to determine how pathogenesis in the absence of Dnase1L3 arises and its impact on specific lupus phenotypes.

One key unknown remaining is the amount of Dnase1L3 needed to prevent disease. Since partial reduction of Dnase1L3 activity contributes to sporadic lupus [3, 8] and HUVS, we aimed to determine how partial reduction of Dnase1L3 impacts autoimmunity in B6 mice. To test how partial reduction of Dnase1L3 impacts lupus-like phenotypes, we generated a Dnase1L3 cell-specific knockout mouse model. In this model, loss of *Dnase1L3* from macrophages caused mild lupus-like phenotypes. Autoantibodies were generated, including anti-dsDNA antibodies, though onset was delayed compared to global Dnase1L3^−/−^. Kidney damage in these mice was mild. These data reveal the lupus-like phenotypes most affected by partial Dnase1L3 loss, and suggest maintaining healthy serum Dnase1L3 activity is crucial to preventing autoimmunity.

## Results

### Macrophages supply half of serum Dnase1L3

To develop a mouse model with a partial reduction in Dnase1L3, we elected to eliminate Dnase1L3 from macrophages. Macrophages were chosen instead of dendritic cells because Dnase1L3 secretion and activity is better characterized in macrophages [11, 17, 19]. To generate cell specific Dnase1L3 conditional knockout (cKO) mice, loxP sites were introduced flanking exons 3 and exon 4 in B6 embryonic stem cells (Supplementary Fig S1A). The neomycin selection cassette was removed by breeding heterozygotes to FLP deleter mice. Mice homozygous for loxP sites flanking Dnase1L3 (Dnase1L3^fl/fl^) were generated, and then crossed twice to LysM-Cre transgenic mice. Mice were born in the expected Mendelian ratio, with 25% Dnase1L3^fl/fl^ x LysM-Cre^+/−^ (cell-specific knockout (cKO)), 25% Dnase1L3^fl/+^ x LysMz-Cre^+/−^ (heterozygous), 25% Dnase1L3^fl/fl^ x LysM-Cre^−/−^ (wild type (WT)), and 25% Dnase1L3^fl/+^ x LysM-Cre^−/−^ (WT). Genotyping was confirmed by PCR (Supplementary Fig S1B). Thus, we developed macrophage-specific *Dnase1L3* knockout mice.

We first assessed these mice for Dnase1L3 levels and Dnase1 activity. Serum was collected weekly from WT and cKO mice. In order to measure mouse Dnase1L3 from mouse serum, we generated polyclonal antibodies to murine Dnase1L3. Immune serum containing the polyclonal antibodies recognized both murine and human Dnase1L3 by ELISA and western blot (Supplementary Fig S2). Using this antibody, we measured serum Dnase1L3 levels by ELISA. Serum Dnase1L3 levels were 67% reduced in the cKO sera compared to the WT (Fig 1A). These results are consistent with previous results [2] using chlodronate liposomes to transiently deplete macrophages. Thus, depletion of Dnase1L3 from macrophages reduces serum Dnase1L3 levels.

**Figure 1.**
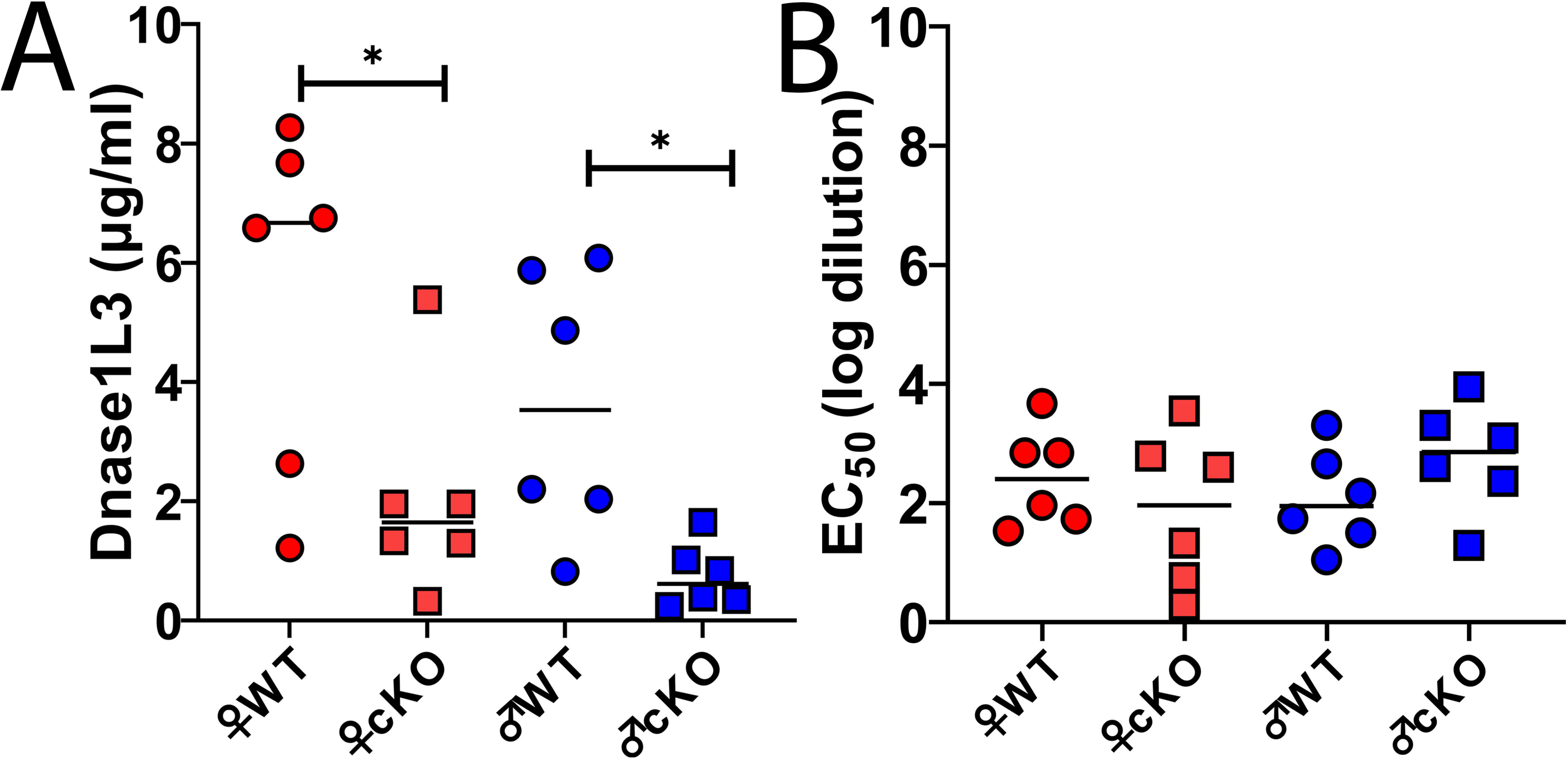
Dnase1L3 loss from macrophages reduces serum Dnase1L3. (A) Serum Dnase1L3 was measured in 41-50 week old Dnase1L3^fl/fl^ x Cre^−/−^ (WT) or Dnase1L3^fl/fl^ x Cre ^+/−^ (cKO) mice by ELISA. (B) Serum Dnase1 activity for 41-50 week old WT or cKO mice was measured using 200 ng plasmid DNA for 30 min. The extent of plasmid degradation was quantified and EC_50_ calculated as described in the methods. Data points represent individual mice and median. Six mice per group were used. * p < 0.05 by one-way ANOVA and Tukey’s multiple comparison test.

We next measured serum Dnase1 activity, which is the ability of nucleases to degrade naked DNA. Dnase1 activity was measured by plasmid degradation, which can be accomplished by both Dnase1 and Dnase1L3 present in the serum [2, 15, 16]. We observed no difference in the Dnase1 activity between WT and cKO mouse sera (Fig 1B). We were unable to measure Dnase1L3-specific activity with either our immune complex degradation assay or barrier to transfection assay [15] because confounders in the serum interfered with these assays. Overall, these data indicate that total serum Dnase1 activity was not altered by macrophage-specific loss of Dnase1L3.

### Loss of Dnase1L3 from macrophages elevates total IgG and IgM antibody levels

We next compared serologic lupus-like phenotypes between WT and cKO mice. To monitor disease progression and construct a timeframe of onset, we collected serum from the mice weekly over their life span. To measure overall induction of autoantibodies, we measured total IgG and IgM levels. Neither antibody subtype was elevated in the first 30 weeks. At 31-40 weeks, we observed an increase in total IgM levels (Fig 2), but no increase in total IgG (Fig 3). By 41-50 weeks of age, both total IgG and IgM levels were elevated 2-fold in cKO mice compared to WT mice (Figs 2A, 3A). A comparison of antibody levels over time revealed a steady increase in both IgM (Fig 2B) and IgG (Fig 3B) for cKO mice. We did not observe any sex-specific differences in elevation of total IgM (Fig 2) or total IgG (Fig 3). These data suggest that reduced levels of Dnase1L3 in macrophages causes late elevation of autoantibodies.

**Figure 2.**
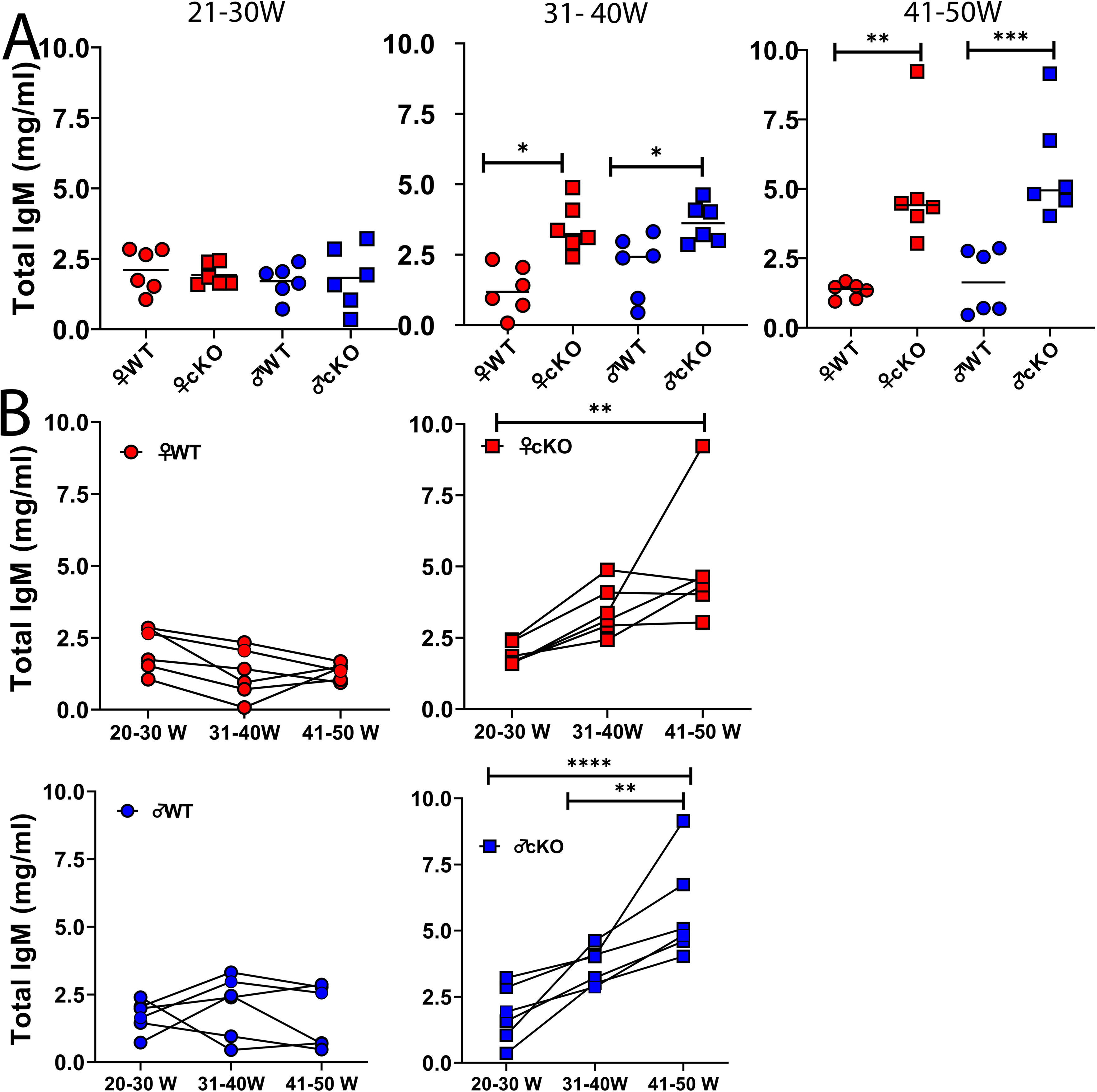
Macrophage-specific loss of Dnase1L3 causes late elevation of serum IgM titers. ELISA for IgM was performed on serially sampled sera from Dnase1L3^fl/fl^ x Cre^−/−^ (WT) or Dnase1L3^fl/fl^ x Cre^+/−^ (cKO) mice between 21-30, 31-40 and 41-50 weeks. (A) Serum IgM levels at each time point are shown. (B) Progression of serum IgM levels with age for each mouse is shown. Graphs show individual mice and median. Six mice per group were used. ***p < 0.001, **p < 0.01, *p < 0.05 by one-way ANOVA, mixed effect models and multiple comparisons using Tukey.

**Figure 3.**
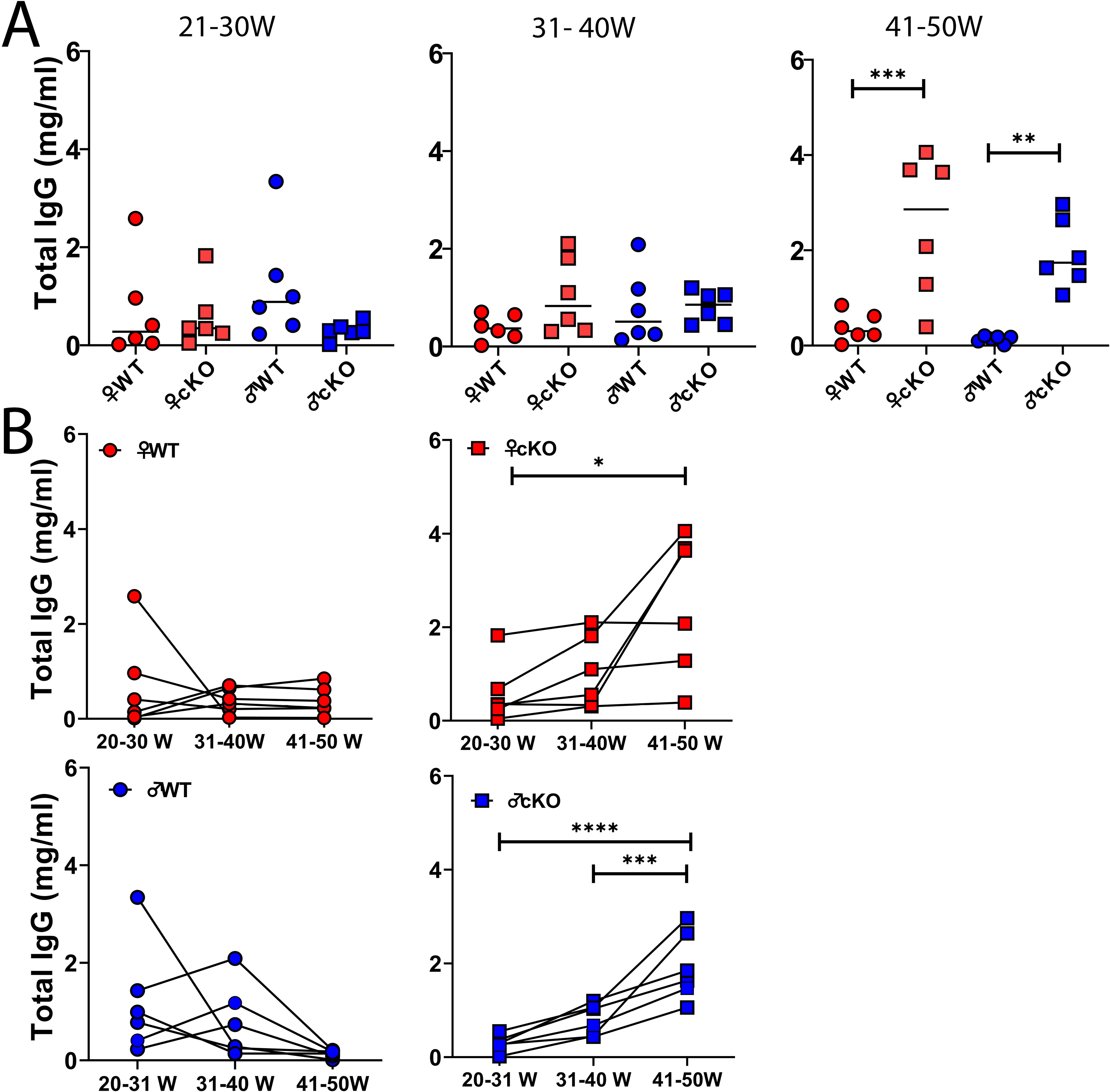
Macrophage-specific loss of Dnase1L3 causes late elevation of serum IgG titers. ELISA for IgG was performed on serially sampled sera from Dnase1L3^fl/fl^ x Cre^−/−^ (WT) or Dnase1L3^fl/fl^ x Cre^+/−^ (cKO) mice between 21-30, 31-40 and 41-50 weeks. (A) Serum IgG levels at each time point are shown. (B) Progression of serum IgG levels with age for each mouse is shown. Graphs show individual mice and median. Six mice per group were used. ***p < 0.001, **p < 0.01, *p < 0.05 by one-way ANOVA, mixed effect models and multiple comparisons using Tukey.

### Partial loss of Dnase1L3 delays onset of anti-dsDNA antibodies

Since total antibody levels were increased in cKO mice, we tested if autoantibodies were produced. We measured anti-nuclear antibody (ANA) titers using commercial HEp-2 cell preparations. In contrast to WT mice, cKO mice had positive ANA titers at the highest dilutions tested, 1:640 (Fig 4). In the cKO mice, the ANA staining pattern was homogenous and perinuclear (Fig 4A). These staining patterns are characteristic of SLE, typically associated with anti-DNA autoantibodies. We next compared anti-DNA antibodies between the serum of WT and cKO mice. Anti-dsDNA antibodies in the serum were elevated at 31-40 weeks (Fig 5A). Female mice showed a greater difference in anti-dsDNA levels compared to male mice (Fig 5A). When examined longitudinally, both male and female cKO showed significant increases in anti-dsDNA antibody levels with age (Fig 5B). These results contrast with a prior analysis of global Dnase1L3^−/−^ mice [2], which had elevated anti-dsDNA antibodies starting at 4 weeks of age. Overall, partial loss of Dnase1L3 was sufficient to generate anti-dsDNA antibodies.

**Figure 4.**
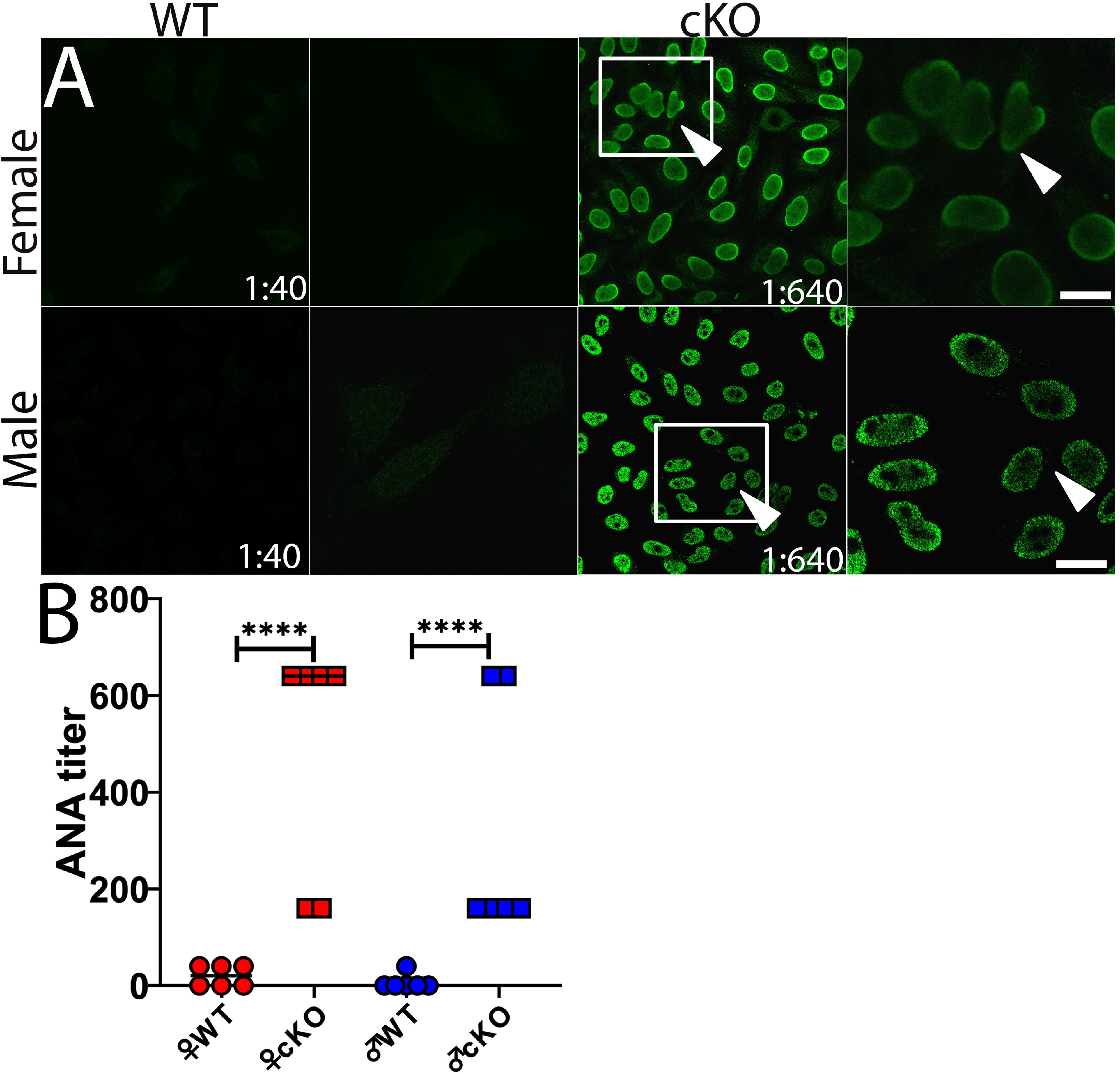
Mice lacking macrophage-specific Dnase1L3 produce anti-nuclear antibodies. Fixed HEp2 cells were incubated with sera from 41-50-week mice at, 1:40, 1:160 or 1:640 dilutions, and stained with DAPI (blue) and anti-mouse IgG (green). (A) Micrographs show a representative field at the indicated dilution. Scale bar = 50 μM (B) Quantitation of ANA titer. Graph shows individual mice plus median. Six mice per group were used. ****p < 0.0001 by one-way ANOVA, mixed effect models and multiple comparisons using Tukey.

**Figure 5.**
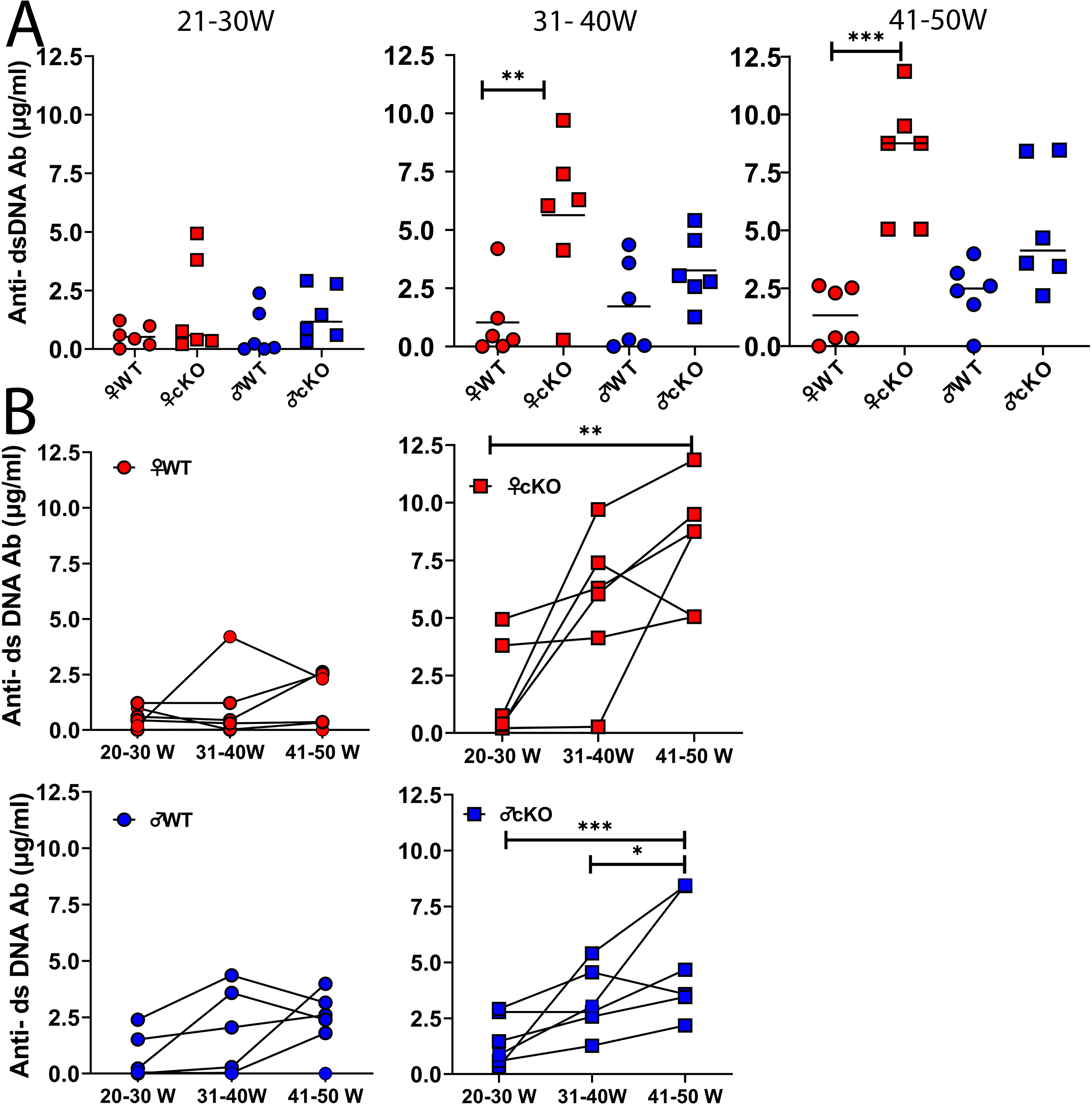
**Elevated anti-nuclear antibodies in macrophage-specific Dnase1L3 knockout mice include anti-dsDNA**. ELISA for anti-dsDNA was performed on serially sampled sera from Dnase1L3^fl/fl^ x Cre^−/−^ (WT) or Dnase1L3^fl/fl^ x Cre^+/−^ (cKO) mice between 21-30, 31-40 and 41-50 weeks. (A) Anti-dsDNA levels at each time point are shown. (B) Progression of anti-dsDNA levels with age for each mouse is shown. Graphs show individual mice and median. Six mice per group were used. ***p < 0.001, **p < 0.01, *p < 0.05 by one-way ANOVA, mixed effect models and multiple comparisons using Tukey.

### Dnase1L3 cKO mice have mild kidney phenotypes

We next compared kidney pathology between the cKO and WT littermate control mice. We euthanized the mice at 50 weeks of age and analyzed their kidneys. Glomeruli had an increased diameter, due to increased thickness of the Bowman’s space in both male and female mice (Fig 6A-C). Kidney pathology scores were elevated, especially in female cKO mice (Fig 6D). Mesangiopathy, hypercellularity, and segmental glomerulonephritis were observed in a subset of the cKO mice, with a greater proportion of female cKO mice showing phenotypes compared to male cKO mice (Fig 6D). This suggests that partial loss of Dnase1L3 does not drive robust kidney pathology.

**Figure 6.**
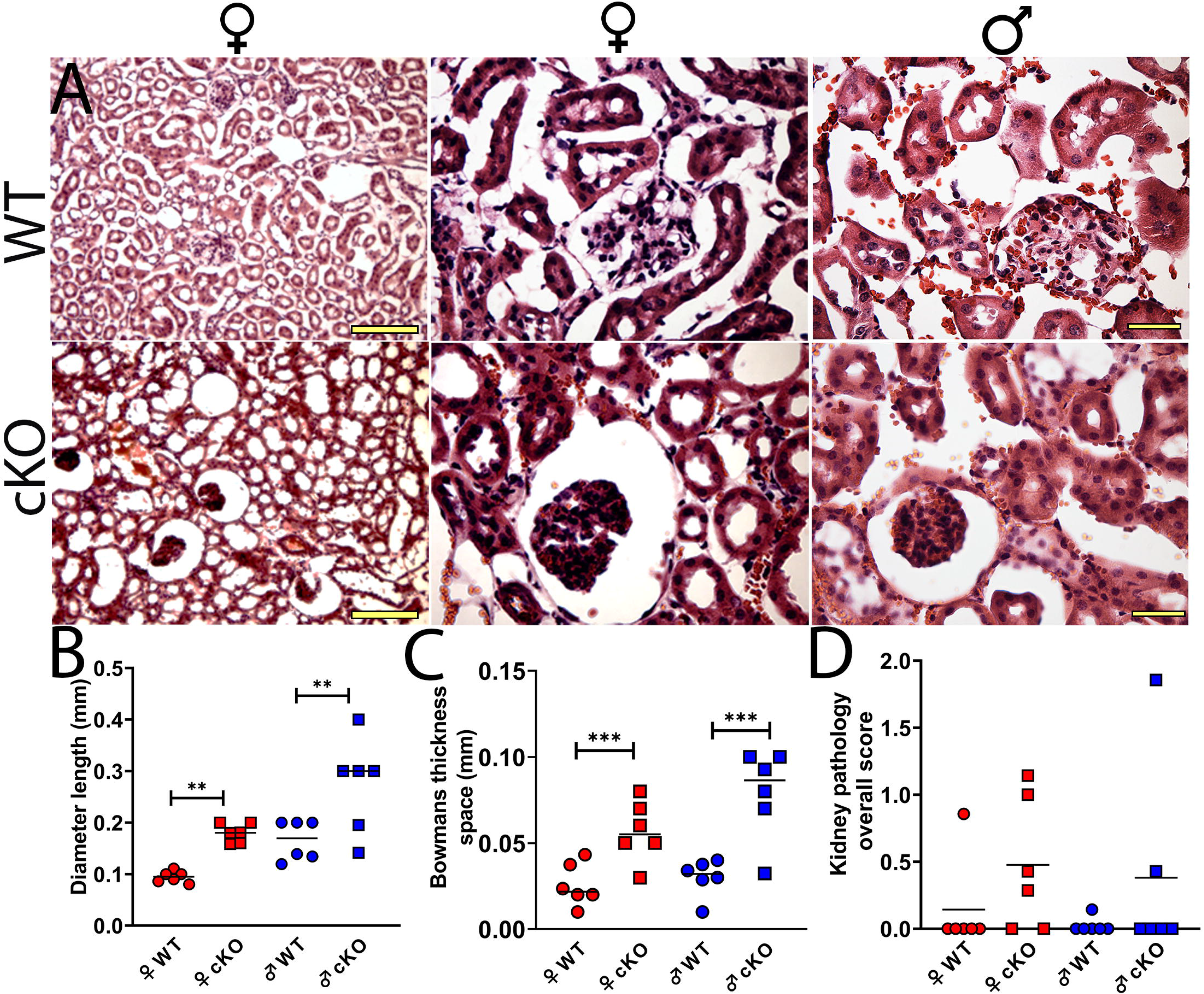
Dnase1L3 loss from macrophages causes mild kidney phenotypes. (A) Kidney sections from 50-week-old Dnase1L3^fl/fl^ x Cre^−/−^ (WT) or Dnase1L3^fl/fl^ x Cre^+/−^ (cKO) mice were stained with hematoxylin and eosin. A representative glomerulus in the inset is shown. (B-C) Glomerular diameter and Bowman’s capsule space were quantitated. (D) The degree of kidney pathology was measured for each mouse. (A) Representative micrographs from six animals per group are shown. (B-D) Graphs show six individual mice per group plus the median for each phenotype. Scale bar = 50 µm *** p < 0.005, ** p < 0.01 by one-way ANOVA and Tukey’s multiple comparison test.

We next measured immune complex deposition in the kidneys. We examined kidney sections for the deposition of IgG, IgM and C3 by immunofluorescence. IgG, IgM and C3 deposited in the glomeruli of cKO, but not WT mice (Fig 7). We observed fine granular immune-complexes and complement deposition in and around the glomeruli (Fig 7A). We quantitated immune complex deposition by measuring the integrated intensities of the glomeruli. IgG, IgM and C3 were elevated in both male and female cKO mice (Fig 7B). These data suggest that partial loss of Dnase1L3 contributes to mild kidney lupus-like phenotypes.

**Figure 7.**
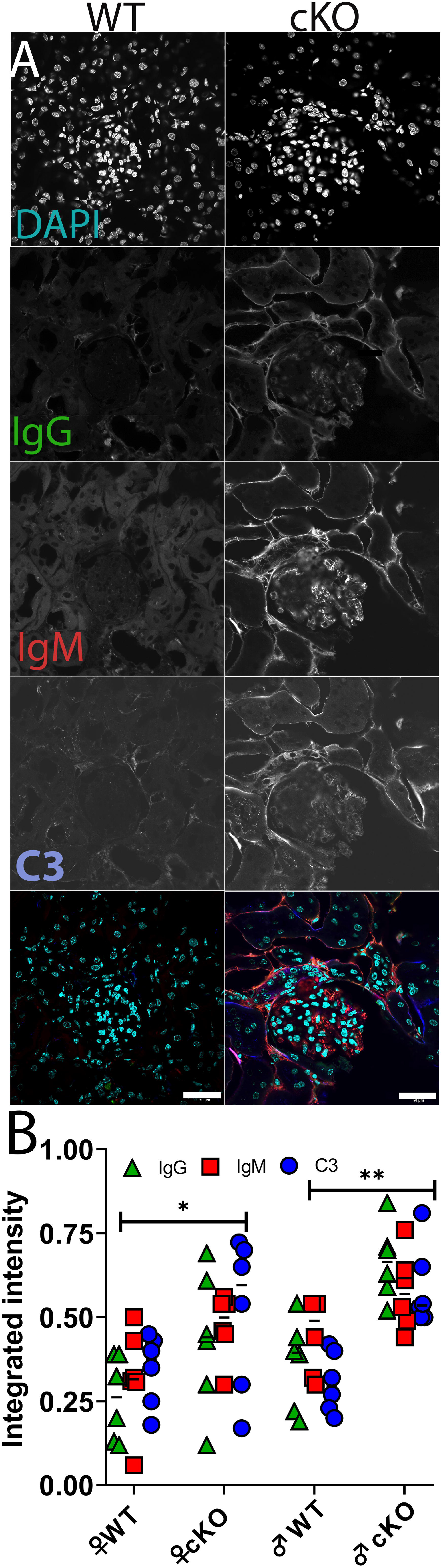
Immune complexes deposit in the kidneys of macrophage-specific Dnase1L3 knockout mice. Kidney sections from 50-week old Dnase1L3^fl/fl^ x Cre^−/−^ (WT) or Dnase1L3^fl/fl^ x Cre^+/−^ (cKO) mice were stained with anti-mouse IgG Alexa 488 (green), DAPI (cyan), anti-mouse IgM Cy3 (red), and anti-C3 Alexa 647 (blue), and imaged by confocal microscopy.(A) representative micrographs or (B) integrated intensities of individual mice with median are shown. Six mice per group were used. Scale bar = 50 µm * p <0.05, ** p < 0.01 by two way ANOVA and Tukey’s multiple comparisons test.

Finally, we compared the lupus-like phenotypes we measured across mice to determine any trends in these phenotypes. We scaled the phenotypes to a 5 point scale, with 5 the maximal phenotype we observed, and 0 the minimum phenotype we observed. Using this scale, we observed an inverse relationship between Dnase1L3 levels and the serology and kidney phenotypes (Fig 8). Due to the mildness of the kidney phenotypes, smaller differences were observed between cKO and WT mice. Overall, we conclude that partial loss of Dnase1L3 is sufficient to trigger mild lupus-like phenotypes in mice.

**Figure 8.**
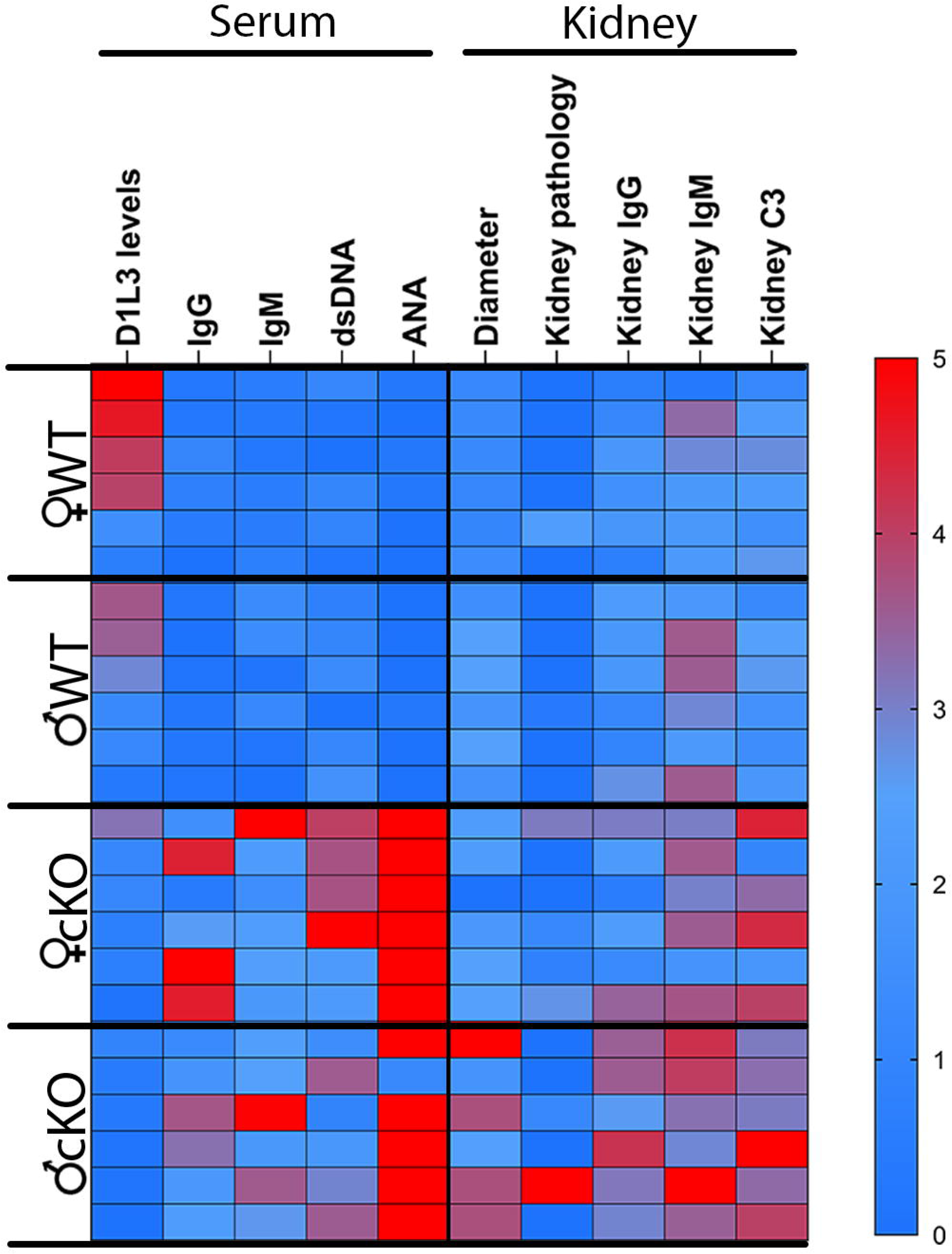
Reduction in Dnase1L3 levels associates with lupus-like phenotypes in mice. Dnase1L3 levels and lupus-like phenotypes were each normalized to a 5-point linear scale, where 5 represents the phenotype of the most affected mouse, and 0 represents the least affected mouse. Each row represents an individual Dnase1L3^fl/fl^ x Cre^−/−^ (WT) or Dnase1L3^fl/fl^ x Cre^+/−^ (cKO) mouse.

## Discussion

In this study, we developed a genetic model of partial Dnase1L3 deficiency by eliminating macrophage-specific *Dnase1L3*. We then used these mice to evaluate the impact of partial Dnase1L3 deficiency on the development of lupus-like phenotypes in mice. While deletion of *Dnase1L3* from macrophages reduced serum Dnase1L3, it did not reduce Dnase1 activity. We found that partial loss of Dnase1L3 increased total IgM, total IgG and autoantibody levels. It caused a delayed increase in anti-dsDNA antibody production relative to full Dnase1L3^−/−^ mice. In contrast, kidney damage was minimal. These results suggest that a partial loss of Dnase1L3 is sufficient to initiate lupus-like phenotypes in mice. Thus, restoring Dnase1L3 may be one promising approach to treat SLE.

While macrophage-specific depletion of Dnase1L3 reduced serum Dnase1L3 50%, it did not alter the serum Dnase1 activity. Serum Dnase1 activity is provided by both Dnase1 and Dnase1L3. One reason for the lack of impact on Dnase1 activity could be a much lower abundance of Dnase1L3 in the serum. Alternatively, compensatory expression of Dnase1 could “correct” for the reduction in Dnase1L3 activity at the level of bulk DNA degradation. For example, polygenic lupus mouse models may attempt to compensate for decreased Dnase1L3 activity by elevating Dnase1L3 expression [17]. However, in our system, production of Dnase1L3 by DCs did not compensate for reduced macrophage expression. Similarly, Dnase1 may compensate for Dnase1L3 activity under certain circumstances [20]. It is possible that Dnase1 compensated for the overall Dnase1 activity. However, Dnase1 cannot rescue many Dnase1L3-specific activities, such as generating extracellular circular chromosomal DNA [21], clearing multi-nucleosome cell-free DNA from mice [22] and humans [23], and preventing lupus onset [2, 14]. Taken together, this suggests any compensatory feedback mechanisms are inadequate to prevent onset of autoimmunity.

Our work provides new insight into lupus heterogeneity and disease onset. Global Dnase1L3^−/−^ mice produce anti-dsDNA antibodies early in life [2, 10, 14]. There is expansion of B cells and neutrophils in the first 8-14 weeks [10], but the full repertoire of autoantibodies do not develop until ∼50 weeks [2]. In contrast, partial loss of Dnase1L3 delayed onset of anti-dsDNA antibodies until 40-50 weeks. We observed the elevation of IgM first, followed by IgG. Interestingly, anti-dsDNA IgG was elevated along with IgM levels, prior to the global increase in IgG. This suggests a gradual accumulation of antigenic extracellular DNA could underlie the trigger for lupus-like phenotypes. The delayed onset of autoantibodies in our model of partial Dnase1L3 loss is more similar to human lupus, where onset often occurs in early adulthood [24]. Importantly, neutralization of Dnase1L3 by autoantibodies occurs in ∼50% of sporadic lupus [8]. The potential for cross-reactivity between anti-DNA and anti-Dnase1L3 [9] autoantibodies suggests a negative feedback loop that accelerates disease. Our work suggests that the partial presence of Dnase1L3 delays this onset. Our macrophage-specific Dnase1L3 knockout represents a genetic system to further explore how partial loss of Dnase1L3 affects the development of autoantibodies. Future work is needed to determine antibody specificity during disease.

Along with insight into DNA-mediated autoantibody onset, we provide additional evidence on organ-system specific lupus defects. Kidney phenotypes from mice with loss of macrophage-specific Dnase1L3 were mild. mesangiopathy, hypercellularity, and segmental glomerulonephritis more typically associated with lupus was mild and present only in a subset of mice. This mild kidney phenotype suggests that loss of macrophage-derived Dnase1L3 is not a major driver of kidney pathology. While the late onset of autoantibodies may have delayed the kidney pathology, similar results from the global knockouts [2] suggest that Dnase1L3 is not uniquely necessary to prevent lupus nephritis in mice. In these circumstances, Dnase1 may rescue kidney phenotypes. For example, Dnase1 rescues renal pathology in *Staphylococcus aureus* infection of Dnase1^−/−^/Dnase1L3^−/−^ mice [14]. Overall, these results provide new insight into our understanding of lupus disease heterogeneity.

Our work opens new avenues of research in the field of lupus and nucleases. For example, it is not clear if dendritic cell-specific deficiency of Dnase1L3 would phenocopy macrophage-specific deletion of Dnase1L3. Our findings suggest that restoration of Dnase1L3 is a viable therapeutic strategy for SLE. Further research is needed to establish the therapeutic potential of Dnase1L3 in SLE treatment.

While we generated and characterized a mouse model with partial Dnase1L3 loss, our work had limitations. Our assessment of lupus-like phenotypes was restricted to serology and kidney pathology. We did not evaluate T cell or B cell subsets nor behavior. Besides anti-dsDNA autoantibodies, we did not characterize specific autoantibody subsets. In global Dnase1L3^−/−^, these repertoires were established ∼40-50 weeks of age [2]. While Dnase1L3 is elevated in M2a macrophages [11], we did not measure extent of macrophage polarization in Dnase1L3 deficient macrophages in vivo. Finally, we did not test the intracellular roles for Dnase1L3 [19, 21].

Overall, our study highlights the importance of optimal levels of Dnase1L3 in preventing the development of SLE. Our macrophage-specific Dnase1L3 knockout mouse provides a model in which partial loss of Dnase1L3 can be modeled. This model expands the repertoire and physiology of lupus mouse models, which provides insights into the impact of variable levels of Dnase1L3 on lupus-like disease heterogeneity in mice.

## Materials and Methods

### Reagents

All reagents were from Thermo Fisher Scientific (Waltham, MA) unless otherwise specified. Gel red was from Biotium (Fremont, CA). Taq DNA polymerase was from Syd Labs (Hopkinton, MA). The anti-human Dnase1L3 rabbit polyclonal antibodies were obtained from Abnova (Cat#H00001776-W01P) and Genetex (Cat #GTX114363). The anti-dsDNA monoclonal antibody clone autoanti-dsDNA was deposited to the Developmental Studies Hybridoma Bank (DSHB) by E.W. Voss. It was obtained from the DSHB created by the NICHD of the NIH and maintained at the University of Iowa, Department of Biology (Iowa City, IA, USA). Rat anti-mouse C3 was from NOVUS Biologicals (Centennial, CO, USA) (Cat #NB200-540). Mouse IgG (Cat # 015-000-002), anti-mouse IgG (Cat# 115-005-003), and anti-mouse IgM (Cat #715-005-020) antibodies were obtained from Jackson Immunoresearch (West Grove, Pennsylvania).

Mouse IgM (Cat #MGM00) was from Invitrogen. HRP-conjugated anti-mouse IgG (Cat #115-035-146) and anti-mouse IgM (Cat #115-035-075) antibodies were obtained from Jackson Immunoresearch (West Grove, Pennsylvania). Anti-mouse IgM conjugated to Cy3 was from Jackson Immunoresearch (West Grove, Pennsylvania) (Cat #115-165-075), anti-rat IgG conjugated to AlexaFluor647 and anti-mouse IgG conjugated to AlexaFluor488 were from Invitrogen. Commercial HEp-2 cells and ANA staining reagents were from MBL Bion (Des Plaines, IL). GFP-Lamp1 cloned in pCS2+ was used for Dnase1 assays, as previously described [15]. Mouse Dnase1L3 lacking the signal sequence was cloned into p202 using BamHI and XhoI restriction sites. Recombinant human Dnase1L3 in p202 was previously described [15]. Recombinant proteins were purified as previously described [15].

### Dnase1L3 antibody

Mouse Dnase1L3 was purified as previously described [15] and concentrated to 5 mg/mL at 95% purity. Two New Zealand White rabbits were immunized and bled by Pacific Immunology (Ramona, Ca, USA), as overseen by their Institutional Animal Care and Use Committee. After bleeding to collect pre-immune serum, rabbits were immunized once in complete Freund’s adjuvant, a second time on day 21 in incomplete Freund’s adjuvant, a third time on day 42 in incomplete Freund’s adjuvant, and a fourth time on day 70 in incomplete Freund’s adjuvant. Antisera was collected on days 49, 63, 77, 91, with final exsanguination bleeds on days 98 and 101. Antibody validation is summarized in Supplementary Fig S2.

### Animals

B6.129P2-Lyz2^tm1(cre)Ifo^/J (LysM-Cre) mice (Strain #004781) were obtained from The Jackson Laboratory (Bar Harbor, ME) and housed at Texas Tech University under the oversight of the Texas Tech University IACUC. Dnase1L3^fl/fl^ mice were generated by inGenious Targeting Laboratory (Ronkonkoma, NY). A loxP site was introduced upstream of exon 3, which encodes the active site of the Dnase1L3 gene, while a neomycin resistance cassette flanked by FRT sites and a second loxP site was inserted downstream of exon 4 (Supplementary Fig S1A).

After removal of the neomycin cassette by breeding to FLP deleter mice, heterozygotes were crossed to generate mice with two floxxed alleles (Dnase1L3^fl/fl^). The breeding scheme to generate macrophage-specific Dnase1L3 knockouts involved crossing the Dnase1L3^fl/fl^ mice with LysM-Cre transgenic mice that express Cre recombinase specifically in myeloid cells including macrophages. This caused macrophage-specific deletion of Dnase1L3 in the offspring. Mice were born with the expected Mendelian ratio, 25% Dnase1L3^fl/fl^ x LysM-Cre^+/−^ (cKO), 25% Dnase1L3^fl/+^ x LysM-Cre^+/−^ (heterozygous) and 50% Dnase1L3^fl/fl^ x LysM-Cre^−/−^ (wild type). The cKO mice were genotyped by PCR using primers specific to the Cre and Dnase1L3.The sample size for each experiment was determined based on a power analysis using estimated effect sizes. No randomization was necessary, so this method was not employed in the study. For blinding, mice were numbered and the experimentalist was blinded to the mouse genotype during the assay. Kidney pathology was read in a blinded fashion by a board-certified pathologist (Tanya LeRoith).

### Genotyping

Genomic DNA from tissue biopsies (tail snips or ear punches) was isolated using a Chelex method [25], or an optimized Proteinase K method [26]. For the Chelex method, tissue samples were ground into a uniform suspension in nuclease-free water and mixed with autoclaved 1% saponin (Sigma) in 1x PBS. After a 20 min room temperature incubation, samples were centrifuged at 17,000xg for 2 min. The pellet was washed once in 1x PBS and vortexed in sterile nuclease-free water. Chelex 100 sodium resin (Bio-rad) was added to 4% (w/v) final concentration. Samples were heated at 95°C for 12 min, and then centrifuged at 17,000xg for 2 min. The supernatant containing genomic DNA was used for downstream analysis. For the optimized Proteinase K method, tissue samples were incubated overnight in lysis buffer containing Proteinase K (100 mM Tris, 5 mM EDTA, 0.2% SDS, 200 mM NaCl, 0.4 mg/mL Proteinase K (Sigma), pH 8.5) at 55°C. The next day, ethanol was added to 70% final volume, and samples centrifuged at 16,000xg for 30 min. The pellet was washed three times with 70% ethanol and centrifuged at 16,000xg for 20 min each time. After the final wash, the genomic DNA was resuspended in 1x TE buffer (100 mM Tris, pH 5.0, 1 mM EDTA, 10 mM NaCl) and stored at −20°C.

PCR genotyping was performed using primers from IDT (Coralville, IA). Primers used for Dnase1L3^fl/fl^ mice were WT1 (5’-GGG CTG GCA TAG AGC ATC AT-3’) and SC1 (5’-CAA CTT GAT GTG AAA GGT GGT AGT G-3’). Primers used for LysM-Cre were oIMR3066 (5’-CCC AGA AAT GCC AGA TTA CG-3’), oIMR3067 (5’-CTT GGG CTG CCA GAA TTT CTC-3’), and oIMR3068 (5’-TTA CAG TCG GCC AGG CTG AC-3’). PCR was performed using the following algorithm: an initial denaturation step at 94°C for 5 min, followed by 34 cycles of denaturation at 94°C for 30 sec, annealing at 56°C for 30 sec, and extension at 72°C for 30 sec, followed by a final extension step at 72°C for 5 min. PCR products were analyzed on a 2% agarose gel. The LysM-Cre band was 700 bp, with 350 bp band for wild type. The floxxed Dnase1L3 allele was detected at 525 bp, while the wild type allele was detected at 476 bp.

### Dnase1 Assay

Dnase1 assays were performed as described [15, 19]. Briefly, 200 ng of plasmid DNA was incubated with varying concentrations of mice sera alone in Dnase assay buffer (20 mM Tris, pH 7.4, 5 mM MgCl_2_, 2 mM CaCl_2_) for 30 min at 37_°_C. The extent of DNA degradation was quantitated by measuring the integrated intensity of degraded and intact plasmid DNA from Gel Red-stained agarose gels using Photoshop Creative Suite (Adobe, San Jose, CA) and determining the percent degradation. The EC_50_ for plasmid degradation was calculated from the dose-response curve using regression on the linear portion of the curve.

### ELISA

Total IgG and IgM levels from mouse sera, were determined as described [27]. A 96-well plate was coated with goat anti-mouse IgG or goat anti-mouse IgM capture antibody overnight at 4°C. Wells were washed three times in 1x PBS with 0.05% Tween 20 (PBST), and blocked with 1% BSA in PBST for 2 h at room temperature. Mouse IgG or mouse IgM was used as standards, while mouse sera samples were diluted 1:5000 in 1% BSA in PBST. After 2 h, plates were washed 3x in PBST, incubated with HRP conjugated goat anti-mouse IgG or IgM antibody at 1:20,000 and 1:10,000 respectively, and developed using 0.2 mg/ml TMB (Sigma), 0.015% H_2_O_2_ in 100 mM sodium acetate, pH 5.5. The reaction was stopped with 0.5 M H_2_SO_4_. A_450_ was measured on a Powerwave Microplate Spectrophotometer running Gen5 Data Analysis Software (Bio-tek, Winooski, VT) and antibody concentration determined.

Anti-dsDNA antibodies were measured as described [27]. The 96 well ELISA plates were precoated with 0.05 mg/mL poly-L-lysine at room temp for 20 min, washed with 1x nuclease-free water, and coated with 5 μg/ml calf thymus DNA (Sigma) overnight at 4° C. After washing 3 x in PBST, plates were blocked for 1 h at room temp with 1% BSA in PBST. Anti-dsDNA (clone autoanti-dsDNA) antibody was used as a standard, starting at 500 pg/mL. Mouse sera were diluted 1:500 in 1% BSA in PBST at RT for 2 h. After 2 h, plates were washed 3x in PBST, incubated with HRP conjugated goat anti-mouse IgG antibody at 1:10,000, and developed using 0.2 mg/ml TMB, 0.015% H_2_O_2_ in 100 mM sodium acetate, pH 5.5. The reaction was stopped with 0.5 M H_2_SO_4_. A_450_ was measured and antibody concentration determined.

To measure the Dnase1L3 levels from mouse serum, 96 well plates were coated with 1:500 of serum diluted in 1X PBS, or with a standard curve starting at 25 ng/ml of recombinant mouse Dnase1L3, overnight at 4° C. After overnight incubation and washing 3 x in PBST, plates were blocked for 1 h at room temp with 1% BSA in PBST. Then anti-mouse Dnase1L3 immune serum was added at 1:5000 for 2 h at room temp. Plates were washed 3x in PBST, incubated with HRP conjugated goat anti-rabbit IgG antibody (1:10,000), and developed using 0.2 mg/ml TMB, 0.015% H_2_O_2_ (Walmart, Fayetteville, AR, USA) in 100 mM sodium acetate, pH 5.5. The reaction was stopped with 0.5 M H_2_SO_4_. A_450_ was measured and the Dnase1L3 concentration determined.

### Histopathology

Kidneys were fixed with 4% paraformaldehyde, dehydrated in 30% sucrose, and embedded in Tissue-Plus™ O.C.T. Compound (Scigen, Paramount, CA), and flash frozen in isopentane followed by liquid nitrogen. Tissues were cut into 5 µm cryo-sections using a Leica CM1950. Sections were stained with H&E. Kidney inflammation was evaluated as previously described [28]. Hypercellularity, mesangial proliferation, necrosis, sclerosis, and tubulointerstitial and perivascular inflammation were scored separately on a scale ranging from 0 (none) to 4 (highest), and the cumulative score was calculated. Images were captured using an Olympus BX41 microscope equipped with a 60x (1.40 NA) objective running QCapture Pro 7 (QImaging, Surrey, BC, Canada) and processed by ImageJ software.

### Immunofluorescence

Immunofluorescence was performed on 5 µm kidney cryo-sections. The sections were fixed for 15 minutes in 2% paraformaldehyde, permeabilized and blocked for 15 minutes in 10% goat serum and 0.05% saponin. The sections were then stained with rat anti-mouse C3 monoclonal antibody for 1 hour, followed by goat anti-mouse IgG conjugated to Alexa 488, goat anti-mouse IgM conjugated to Cy3, and goat anti-rat IgG conjugated to Alexa 647 for 1 hour, and finally DAPI stained. The sections were imaged using a Fluoview 3000 confocal microscope (Olympus, Tokyo, Japan) equipped with a 60x, 1.42 NA oil immersion objective.

The deposition of IgG, IgM immune complexes, and Complement protein 3 was determined by measuring the integrated intensities using ImageJ software. The images were analyzed by splitting each image into RGB channels. All fluorescence integrated density values were normalized based on the area and respective backgrounds. Mean density values of each image were calculated, and data was recorded and graphed using GraphPad Prism. The brightness and contrast of all confocal images were adjusted equally for qualitative purposes.

### Anti-nuclear antibody staining

Anti-nuclear antibodies (ANA) were detected using fixed HEp-2 cells (MBL Bion) as a substrate. Mouse serum was incubated with HEp-2 cells at different dilutions (1:40, 1:160, 1:640), slides washed, and then the slides were stained with a goat anti-mouse IgG conjugated to Alexa 488, followed by staining in DAPI. The stained cells were imaged using a Fluoview 3000 confocal microscope (Olympus, Tokyo, Japan) equipped with a 60x, 1.42 NA oil immersion objective. The images were processed using ImageJ. For qualitative purposes, the brightness and contrast were adjusted equally for all confocal images. At least 30 cells were evaluated per sample. Fields of cells with fluorescence >2 standard deviations over the negative control were considered positive.

### SDS-PAGE and immunoblotting

SDS-PAGE and immunoblotting were performed as described [29]. Briefly, samples were resolved on 10% polyacrylamide gels at 160 V for ∼60 min and transferred to nitrocellulose in an ice bath with transfer buffer (15.6 mM Tris and 120 mM glycine) at 300 mA for 90 min. The blots were blocked using 5% skim milk in 10 mM Tris-HCl, pH 7.5, 150 mM NaCl, and 0.1% Tween 20. Portions of the blot were incubated with one of the following primary antibodies for 2 hours at room temperature or overnight: anti-human Dnase1L3 (1:1000) (Abnova and Genetex), pre-immune rabbit serum (1:5000), or rabbit anti-mouse Dnase1L3 immune serum. Next, the blots were washed 3x in TBST, incubated with HRP-conjugated anti-rabbit IgG antibodies (1:10,000), washed 3x in TBST, and developed with enhanced chemiluminescence (ECL): 0.01% H_2_O_2_, 0.2 mM p-Coumaric acid (Sigma-Aldrich), 1.25 mM luminol (Sigma-Aldrich) in 0.1 M Tris (pH 8.4).

### Statistics

Prism 8.1 (GraphPad, San Diego, CA, USA) or Excel were used for statistical analysis. Data are represented as mean ± s.e.m. as indicated. The EC_50_ for Dnase1 activity was calculated in Excel as previously described [30]. Exclusion criteria for data was failure of positive or negative controls. Statistical significance for normally distributed data was determined by one-way or two-way ANOVA with Tukey post-testing as specified; p < 0.05 was considered to be statistically significant. Graphs were generated in Excel, Prism and Photoshop (Adobe, San Jose, CA, USA).

## Acknowledgments

The authors would like to thank members of the Keyel lab for critical review of the manuscript. We thank the College of Arts & Sciences Microscopy for use of facilities. This work was supported by Texas Tech University (PAK), American Heart Association grant 16SDG30200001 (PAK), and Lupus Research Alliance grant 707050 (PAK). CJH was supported by the Texas Tech University, Ronald E. McNair Post-Baccalaureate Achievement Program, U.S. Department of Education Grant, 2017-2022. AI was supported by the Plains Bridges to the Baccalaureate NIH grant R25GM083730.

## Author Contributions

ME: Conceptualization, Investigation, Data curation—Formal analysis, Visualization, Validation, Writing—Original Draft, Writing—Review and Editing

CH: Investigation, Data curation—Formal analysis, Visualization, Writing—Original Draft, Writing—Review and Editing

AI: Investigation, Data curation—Formal analysis, Visualization, Writing—Original Draft, Writing—Review and Editing

TL: Investigation, Data curation—Formal analysis, Writing—Review and Editing

ER: Investigation, Data curation—Formal analysis, Writing—Original Draft, Writing—Review and Editing

LE: Investigation, Data curation—Formal analysis, Writing—Review and Editing JV: Investigation, Resources, Writing—Review and Editing

CR: Validation, Data curation—Formal analysis, Writing—Review and Editing, Project Administration

RBS: Validation, Resources, Writing—Review and Editing, Funding Acquisition, Project Administration

PAK: Conceptualization, Investigation, Validation, Data curation—Formal analysis, Supervision, Writing—Original Draft, Writing—Review and Editing, Funding Acquisition, Project Administration

## Conflicts of Interest

PAK <u>and</u> RBS have patents pending on Dnase1L3 modifications to improve serum half-life. The funders had no role in the design of the study; in the collection, analysis, or interpretation of data; in the writing of the manuscript; nor in the decision to publish the results. The content is solely the responsibility of the authors and does not necessarily represent the official views of the funding agencies.

## Data and materials availability

All data are available in the main text or the supplementary materials.

## Supplementary Figure Legends

**Supplementary Figure S1. Generation of Dnase1L3^fl/fl^ and Dnase1L3^fl/fl^ x LysM-Cre^+/−^ mice.** (A) The schematic illustrating the targeting strategy for generating the Dnase1L3 conditional knockout mouse model. Exons 3 and 4 of *Dnase1L3* were flanked with loxP sites, with a neomycin cassette introduced downstream of exon 4. Transgenic mice with germline integration of the targeting vector were bred to FLP deleter mice to remove the Neomycin cassette (Neo deletion). These mice were used as wild type mice. To generate cell-specific mouse knockouts, Dnase1L3^fl/fl^ mice were bred to LysM-Cre (Dnase1L3 x Cre). (B) Genotyping of mice was performed by PCR. The Dnase1L3 wild-type allele yielded a PCR product of 476 bp, while the mutant allele yielded a PCR product of 525 bp (top). For LysM-Cre, the wild type and mutant alleles are represented by 350 and 700 bp, respectively (bottom).

**Supplementary Figure S2. Validation of the anti-mouse Dnase1L3 polyclonal antibody.** (A-B) Either (A) 20 ng or (B) 5 µg recombinant human (Hs D1L3) or mouse (Mm D1L3) was resolved by SDS-PAGE and (A) transferred to nitrocellulose for western blotting, or (B) Coomassie stained to show protein loading. Membranes were probed with pre-immune sera, sera from two rabbits immunized against mouse Dnase1L3, or two commercially available anti-human Dnase1L3 antibodies (Abnova and Genetex), followed by anti-rabbit IgG conjugated to HRP. (C) ELISA was performed by coating plates with recombinant mouse Dnase1L3, and adding increasing concentrations of pre-immune or immune serum. The graph displays the mean ± SEM of three independent experiments.

## Notes

### Competing Interest Statement

PAK and RBS have patents pending on Dnase1L3 modifications to improve serum half-life. The funders had no role in the design of the study; in the collection, analysis, or interpretation of data; in the writing of the manuscript; nor in the decision to publish the results. The content is solely the responsibility of the authors and does not necessarily represent the official views of the funding agencies.

