## Supplementary figures and images for "Deficiency of macrophage-derived Dnase1L3 causes lupus-like phenotypes in mice"

### Supplemental Figures

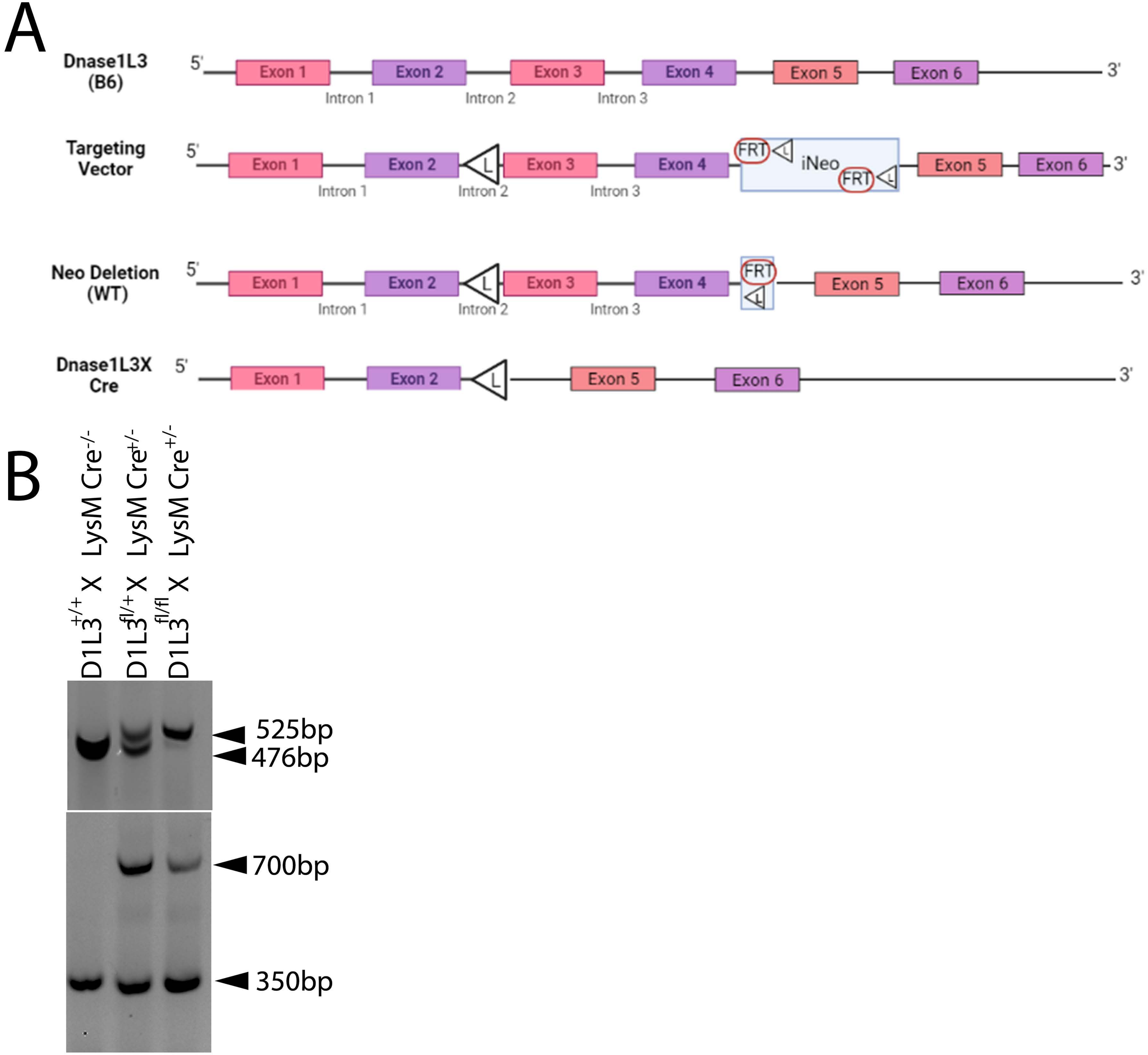

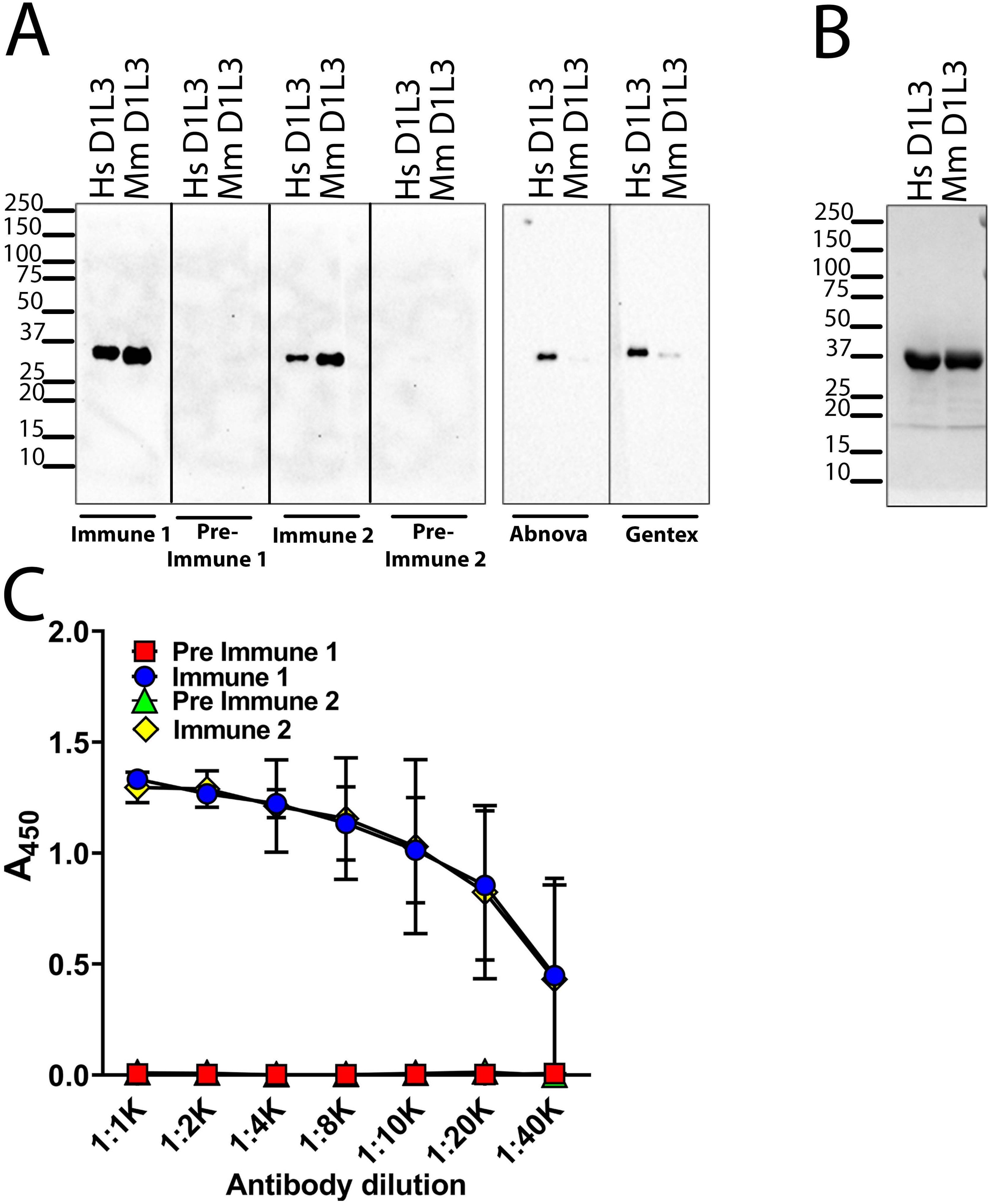
